# Investigating the impact of fibroblast proximity to a tumor on fibroblast extracellular vesicle production utilizing 3D bioprinted stromal models

**DOI:** 10.1101/2024.12.02.626455

**Authors:** Jensen N. Amens, Jun Yang, Lauren Hawthorne, Pinar Zorlutuna

## Abstract

Extracellular vesicles (EVs) are an important carrier of cellular communication that are secreted from the cell. Different cells will produce EVs with different cargo such as cytokines, RNAs, or microRNAs (miRNA). EVs have been proven to play an important role in breast cancer tumorigenesis, progression, and metastasis. Although the role of cancer associated fibroblasts (CAFs), and EVs originated from them have been studied extensively, there is a lack in knowledge on the contribution of normal fibroblasts surrounding the tumor and their roles with respect to their proximity to the tumor. Here we investigate how the proximity of the tumor affects the EV production of the normal fibroblasts. We created stromal models by 3D bioprinting two different fibroblasts, normal human mammary fibroblasts (hMFs) and normal tumor adjacent fibroblasts (NTAF), within a collagen gel. After one week of culture, we isolated EVs from both the effluent media and the 3D stromal model, which were then characterized using nanoparticle tracking analysis (NTA), transmission electron microscopy (TEM), ELISA, zeta potential, and cytokine array analysis of the cargo. The EVs from each group were of consistent exosome size and displayed traditional exosome markers, however the EVs from different groups also displayed different cytokine profiles of their cargo, with the NTAF media group showing an upregulation of cytokines associated with breast cancer progression. After this, we used the EVs to treat breast cancer cells to investigate the effects the EVs from different tumor proximities have on the breast cancer cell behavior. The breast cancer cells treated with the NTAF groups had increased migration. Finally, we utilized a 3D breast tumor model to investigate the effects of the EVs on a tumor spheroid. Tumor spheroids treated with either NTAF EV groups showed increased proliferation, tumor radius, and local invasion. This study is the first to investigate the effect of proximity to a breast tumor on EV production and the first to utilize 3D bioprinting of stromal models specifically to obtain EVs. Overall, our results show that EVs from normal fibroblasts closer to a tumor produce EVs that promote breast cancer progression, regardless of the secretion location of the EVs. These cells have a distinct EV secretome different from normal human mammary fibroblasts, showing that the proximity to a tumor influences the normal fibroblasts surrounding the tumor.

## Introduction

Extracellular vesicles are small particles with a bilipid membrane that are released from the cells^1,2^ which play an important role in intercellular communication. They typically contain various types of RNAs – including microRNAs (miRNA) – cytokines, and other molecules used for cell signaling with different types of cells producing different cargo. The cargo from EVs can affect various cellular and organ processes with EVs from dysfunctional cells carrying pathogenic proteins or proinflammatory cytokines to progress various diseases such as renal disease ^3^, neurodegenerative diseases ^4,5^, inflammatory diseases ^6^, and various types of cancers ^7,8^. EVs have been proven to play an important role in breast cancer tumorigenesis, progression, and metastasis ^9–11^ and can contain important biomarkers for breast cancer ^12,13^, which could allow early detection of breast cancer and provide a more accurate prognosis based on content from the EVs from various biological fluids.

While many studies have been focused on EVs produced from biological fluids and cellular secretions, recently, there is growing interest in a novel type of EVs bound within the ECM of tissues. It has been shown that EVs isolated from the ECM of specific tissues contain different contents than those isolated from fluids ^9,14–18^. In one study, cells treated with EVs isolated from ECM had increased neurite outgrowth, proving the EVs from ECM are bioactive like their fluid counterparts ^17^. Similar results have also been obtained for commercially available ECM scaffolds ^14^, cancer associated fibroblast (CAF)-derived ECM ^15^, urinary bladder ECM ^17^, and others. Previous studies within our lab have shown differences between the EVs that are present within aged and young breast tissue ^9^ and discovered that the aged ECM EVs caused breast cancer cells to become more motile and invasive. These aged ECM EVs also contained more cytokines and miRNAs that are associated with cancer initiation, metastasis, and treatment resistance. Multiple studies from our lab ^18,19^ and others^14–17^ have shown, EVs isolated from ECM are an important source to study as they may provide more insight to breast cancer initiation, progression, and metastasis. Along with changes in EVs from different secretion locations, as stated previously, different cell types can also produce differing EVs. Fibroblasts within the breast tumor, CAFs, have been shown to secrete EVs that aid in breast cancer progression and metastasis ^20–22^. EVs from CAFs have been well characterized through transmission electron microscopy (TEM), nanoparticle tracking analysis (NTA), qRT-PCR, western blot, microRNA profiling, and other methods to characterize their size, shape, and cargo; however, EVs from normal fibroblasts surrounding the tumor have not been explored. It is necessary to investigate the changes in EVs from different cell types and secretion locations.

Because EVs can differ based on their place of uptake – such as biofluids, cell culture media, or ECM – it is important to have relevant models when investigating EVs to understand the different properties and functions of EVs with different biogenesis. 3D engineered models have been proven to allow researchers to investigate cell-cell and cell-ECM interactions in a more physiologically relevant environment ^23–26^. These models can be produced using patient-derived tumor cells and ECM to better replicate the specific microenvironment being studied; however, these 3D engineered models have limitations that include reproducibility within models, scaling production of models, and the heterogeneity of the tumor microenvironment ^24^. One method to help solve some of these limitations is to utilize 3D bioprinting, which can overcome reproducibility and production scaling as the bioprinter can simultaneously print multiple models, limiting differences between models and allowing for many models to be printed at once. This method can also recreate the heterogeneity of the tumor microenvironment through printing multiple types of materials, cells, or cytokines in specific locations within the 3D model. 3D bioprinting has been used to print various tissues such as vessels ^27–30^, bone ^31,32^, and cartilage ^33–35^, as well as drug-screening model systems ^36,37^. One of the main application for 3D bioprinting within cancer research is to create 3D cancer models that better represent the human body environment as well as create high throughput systems for cancer research. Previous research has included printing tumors within a stromal gel to create a relevant cancer model ^38,39^ to better aid in drug development while another study printed cancer cells and surrounded them by supporting adipose-derived mesenchymal stem cells (ADMSCs) to investigate drug resistance^40^. Other studies have utilized this biofabrication method to pattern cells in specific places to better mimic the tumor microenvironment ^41^. This method has been shown to improve engineered models and replicate the tumor microenvironment. This could be utilized to investigate EVs within more biomimetic environments to better understand the role EVs play in disease progression. Previous studies combining 3D bioprinting and EVs focus on printing the EVs into specific places ^42–46^ or treating constructs with EVs ^47^ rather than printing a construct and isolating EVs from this construct as we have shown here. 3D bioprinting allowed us to create a stromal model for obtaining EVs in a reproducible and human mimicking manner.

This study aims to investigate how the proximity of the tumor affects the EV production of normal fibroblasts by utilizing 3D models of the breast stroma and the breast tumor microenvironment. To examine this, we created stromal models by 3D bioprinting the two different normal fibroblasts, normal human mammary fibroblasts (hMFs) and normal tumor adjacent fibroblasts (NTAF), within a collagen gel. They were cultured for one week before isolating EVs from both the effluent media and the 3D stromal model. The resulting EVs were characterized using NTA, TEM, ELISA, zeta potential, and cytokine array analysis of the cargo. Through our characterizations we found that the EVs were of consistent exosome size and displayed traditional exosomal markers, while their concentration and surface charge differed based on their origin. The EVs also displayed different cytokine profiles of their cargo, with the NTAF media group showing an upregulation of cytokines associated with breast cancer progression. After this, we used the EVs to treat breast cancer cells to investigate the effects the EVs have on the breast cancer cell behavior. The breast cancer cells within a 2D environment treated with the NTAF groups had increased migration. Finally, we utilized a 3D breast tumor model to investigate the effects of the EVs on a tumor spheroid which caused an increase in the local invasion and proliferation of the 3D tumor spheroid. This study is the first to investigate the effect of proximity to a breast tumor on stromal cell EV production as well as the first study to utilize 3D bioprinting to obtain EVs. While previous studies have characterized CAFs, we focus on a different subpopulation of normal fibroblasts that are adjacent to the tumor rather than within the tumor as CAFs are. Overall, our results show that normal fibroblasts closer to a tumor produce EVs, independent of the EV secretion origin, that promote breast cancer progression.

## Materials and Methods

### Cell Culture

GFP-reporting MDA-MB-231 breast cancer cells were a kind gift from Dr. Siyuan Zhang at University of Notre Dame. Breast cancer cells were cultured in medium (DMEM high glucose medium supplemented with 10% fetal bovine serum (FBS) and 1% penicillin-streptomycin (P/S)). The culture was maintained with media changes every two days until 90% confluent.

hMFs and NTAFs were obtained from Prof. Harikrishna Nakshatri at Indiana University School of Medicine ^48–51^. Primary cells were cultured in basal medium ((DMEM [low glucose]: Ham’s F12 [1:3] medium supplemented with 5% FBS [Thermo Fisher Scientific], 0.4 μL/mL hydrocortisone [Sigma], 1% penicillin/streptomycin [Corning], 5 μg/mL insulin [Sigma], 10 ng/mL EGF [Millipore], 6 mg/mL Adenine [Sigma], and 10 mM ROCK inhibitor [Y-27632] [Enzo Life Sciences]).

### Stromal Model Fabrication

Fibroblasts were detached using trypsin. These were then mixed into a 6.6 mg/mL solution of human collagen (HumaBiologics, USA) at a concentration of 20 million cells/mL, loaded into a bioprinter (Cellink, USA), and droplet printed into a 96-well plate to form stromal models.

Stromal models were printed with pressures from 7-14 kPa and extrusion times from 0.70-0.85 sec. The stromal model systems were allowed to gel at 37 °C before being cultured in basal medium for up to seven days with a media change after three days in culture.

### Extracellular Vesicle Isolation

Model EV isolation was performed based on a modification of a previously described protocol ^52^. Stromal models were decellularized with the decellularization solution and rinsed with PBS and deionized water for rehydration. To release the model EVs from the matrix, stromal models were enzymatically digested with a solution containing 0.1 mg/mL type II collagenase, 50 mM tris buffered saline (TBS), 5 mM CaCl2 and 200 mM NaCl for 24 h at room temperature with agitation. Enzymatically digested models were subjected to successive centrifugations at 500g for 10 min, 2,500g for 20 min and 10,000g for 30 min. Each centrifugation step was performed three times to ensure the removal of collagen fibril remnants. The fiber-free supernatant was then centrifuged at 100,000g with tabletop ultracentrifuge (Beckman Coulter) for 70 min at 4 °C. The pellets were stored at −80 °C or resuspended in PBS for immediate use. Media collected from the models after one week in culture was subjected to centrifugation twice at 10,000g for 30 min before being centrifuged at 100,000g with tabletop ultracentrifuge (Beckman Coulter) for 70 min at 4 °C.

### EV Size Determination

EVs from each group were resuspended in PBS and analyzed with a NanoSight NS3000 instrument (Malvern). Size distribution and particle concentration of MBVs were determined by Brownian motion measurement (n = 5). NanoSight data was analyzed with R software to calculate the number and diameter of the particles after reduction of background noises.

### TEM Imaging of EVs

EV size and morphology characterization was further confirmed with TEM imaging. Briefly, EV samples (n = 3) isolated with ultracentrifugation were resuspended and incubated in 2.5% glutaraldehyde for 1 hour at room temperature and loaded on plasma cleaned carbon-coated copper grid (Polysciences). Fixed EV samples were incubated on the grids for 20 min at room temperature. The samples were washed with deionized water and vacuumed for 10 min to dry.

EV samples were incubated in vanadium solution (Abcam) for 15 min at room temperature for negative staining. The samples were then washed with deionized water and vacuum dried. TEM imaging was performed at 120 kV.

### Surface Marker Characterization

Quantification of surface markers of EVs was performed with ExoELISA-ULTRA Complete Kit for CD9, CD63, and CD81 detection (SBI). Isolated EVs were resuspended in PBS and added into protein-binding 96-well plates at 5 μg per well. Plates were incubated at 37 °C for an hour for protein binding and washed 3 times with 1x wash buffer. Plates were then treated with corresponding primary and horseradish peroxidase-labelled secondary antibodies and super- sensitive tetramethylbenzidine substrates according to manufacturer’s instructions. Reactions were terminated with stop buffer and quantitative results were measured at 450 nm with a plate reader (Wallac 1420).

### Cytokine Profiling

Cytokine profiling of the EV groups was performed using the dot blot-based Proteome Profiler Human XL Cytokine Antibody Array kit (R&D Systems) following the manufacturer’s instructions. EVs were lysed with 1% Triton X-100 overnight at 4 °C. Total protein concentrations were determined by Pierce Gold BCA Assay (Thermo Fisher Scientific), and equal amounts of protein were loaded onto the blots to assess the cytokine levels. Result of the cytokine array was analyzed using ImageJ by quantifying the intensity of the dots from each sample.

### Cell Motility

The effect of the EVs on cell motility was assessed through an *in vitro* migration assay. The MDA-MB-231 cells were seeded in 24-well plates and treated with 20 μg MBVs for 48 h. Cells were then washed, treated with trypsin, and reconstituted in culture media. A 30 μL aliquot of the cell suspension was seeded in the center of a glass bottom culture dish (total of 100,000 cells), incubated overnight at 37°C for the cells to attach. After cell attachment, fresh media was added, and cells were subjected to time lapse imaging under an inverted fluorescence microscope for 10 h with 15 min intervals. Cells were tracked with Fiji software (NIH). Non-treated cells cultured under the same conditions were used as a control.

### Model Fabrication

Tumor spheroids were created by seeding MDA-MB-231 cells into ultra-low attachment plates at a density of 10,000 cells/ μL. These were allowed to culture for two days before droplet printing (Cellink, USA) a 6.6 mg/mL solution of human collagen (HumaBiologics, USA) onto the tumor spheroids. These were allowed to gel at 37 °C prior to placing basal medium onto the constructs and treatment with EVs for two weeks.

### Model Characterization

Viability of cells in models was determined using the live/dead cell viability assay (Thermo Fisher Scientific). Briefly, models (*n* = 3) at day 1 of culture were incubated in a solution of 2 μM calcein AM (live cells, green) and 4 μM ethidium homodimer 1 (EthD 1) (dead cells, red) at 37 °C for 30 min, and cells imaged using a fluorescence microscope.

### Immunostaining

3D tumor models (*n* = 3) were stained for Ki67. Models were permeabilized with 0.1% Triton X 100, and then incubated for one hour in 5% goat serum, overnight at 4 °C in mouse anti-human Ki67 (Abcam) monoclonal antibodies with 1:100 dilution, and then for 6 h at 4 °C in Alexa fluor 647 labelled goat anti mouse IgG (Abcam) with 1:400 dilution. The samples were stained with 0.5 μg/mL DAPI (Sigma) and imaged with a two photon confocal microscope.

### Statistics

Data were analyzed for statistical significance with Prism 9 (Graphpad). One-way ANOVA followed by Tukey’s HSD correction was performed to compare the differences between EV groups. Outliers were identified using the ROUT method with *Q* = 1% and eliminated. Data are presented as the mean ± standard deviation (SD).

## Results

### Creation of the Breast Stromal Model and Cell viability

NTAF cells have been characterized previously^48–51^ and were shown to be a different cell population from CAFs within the breast stroma. Stromal models were created as depicted below in figure 1. Models were stained with calcein AM and ethidium bromide to stain the live cells green and dead cells red. The models showed that after printing the majority of cells were still alive (Figure 2B). hMF models showed an 82.8% viability and NTAF models showed a 76.7% viability.

**Figure 1:**
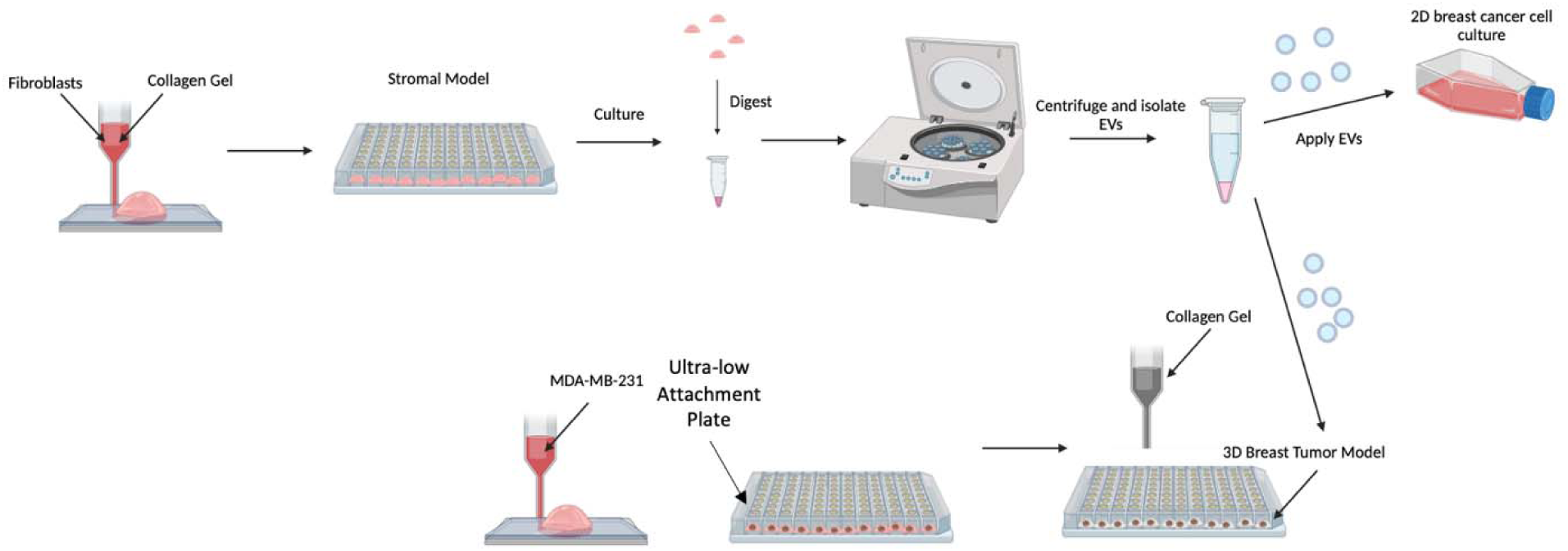
Schematic of stromal model, EV isolation, and cell assay system.

**Figure 2:**
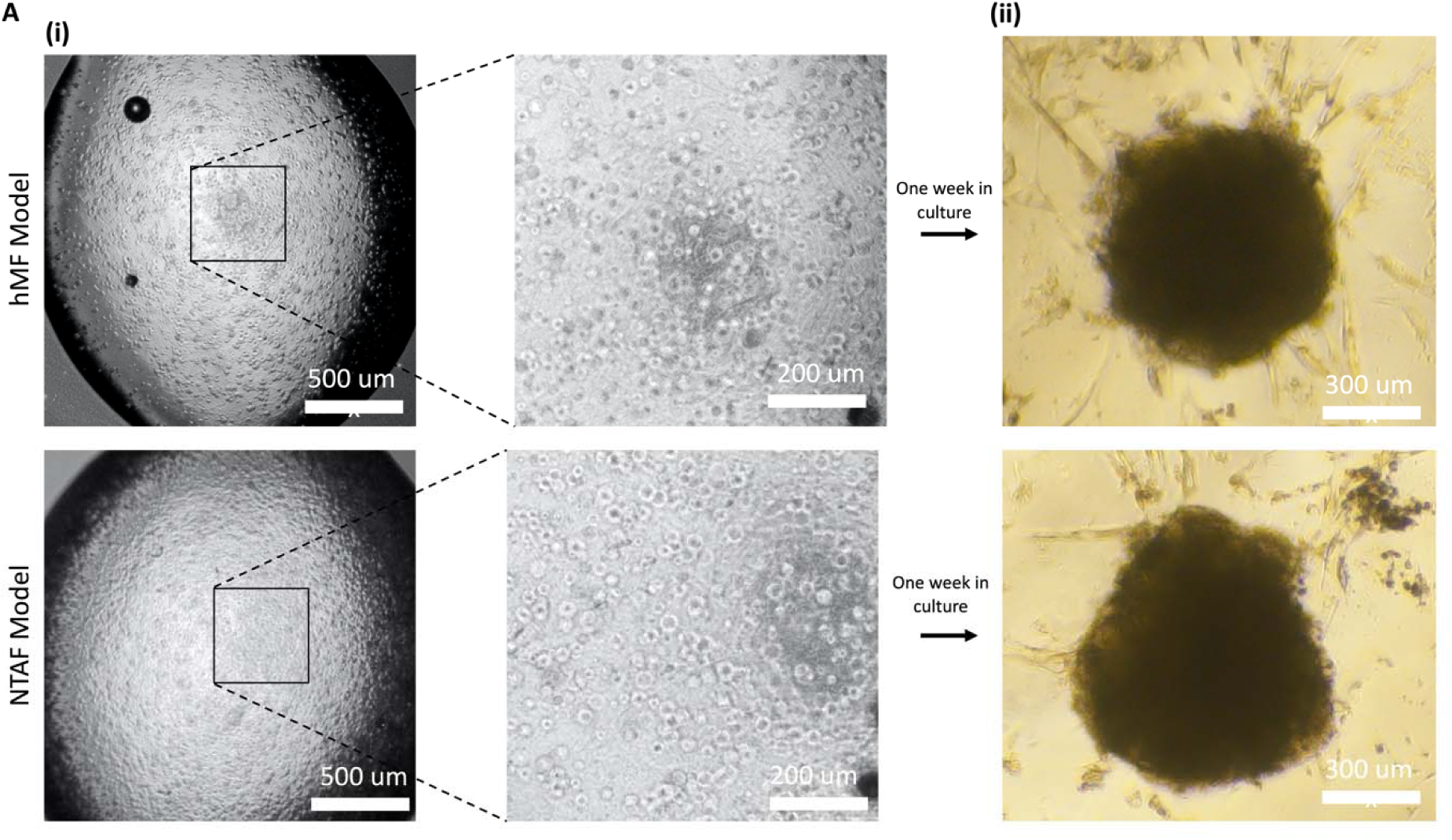
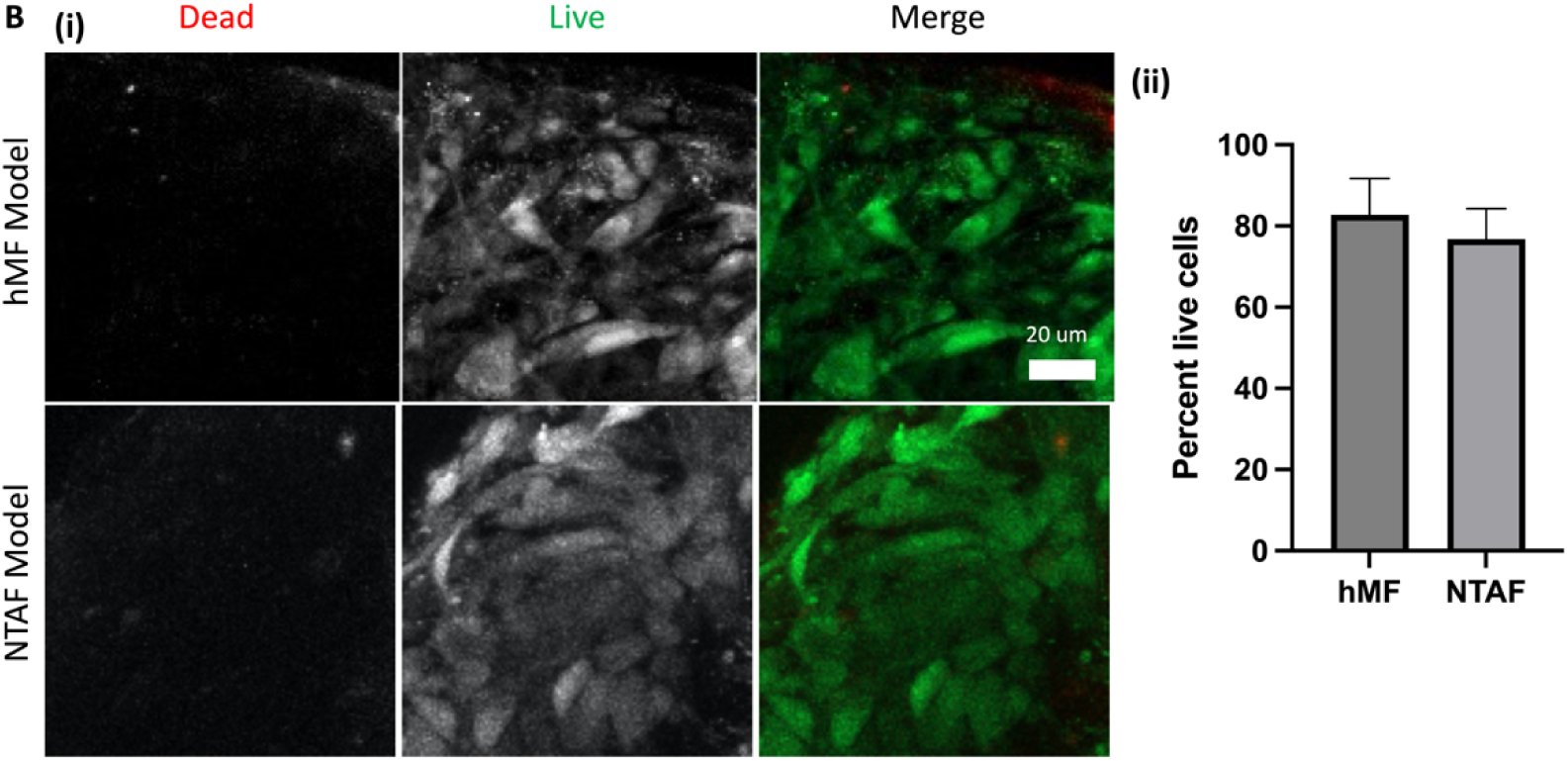
Stromal model images and cell viability. A) (i) Brightfield images of models immediately after printing. (ii) Brightfield images of models after one week in culture. B) (i) Fluorescence images of live/dead stained models. (ii) Quantification of percent live cells in the images (n = 7).

### Stromal model EVs show consistent size and exosome surface markers

Stromal model EVs from both media and within the models were isolated after one week in culture. The pellets were then resuspended in PBS for characterization. First, samples were prepared for TEM. These images show a consistent size and shape throughout the EV groups (Figure 3A). EVs were then characterized by size and concentration using NTA. This confirmed that the EVs are within standard EV size (Figure 3B). This also showed an increase in concentration for the NTAF model EVs as compared to the NTAF media EVs and the reverse for the hMF groups. After this, we characterized common exosome surface markers CD9, CD63, and CD81 through ELISA. We were able to confirm the presence of these surface markers showing that we successfully isolated exosomes (Figure 3C). We saw a difference in the zeta potential, which measures the surface charge of the EVs, between the model groups and the media groups (Figure 3D). hMF media and NTAF media groups had larger average zeta potential at -11.19 and -14.37 respectively. The model groups were closer to zero with the hMF model group having a zeta potential of -1.78 and NTAF model group with a zeta potential of - 1.38.

**Figure 3:**
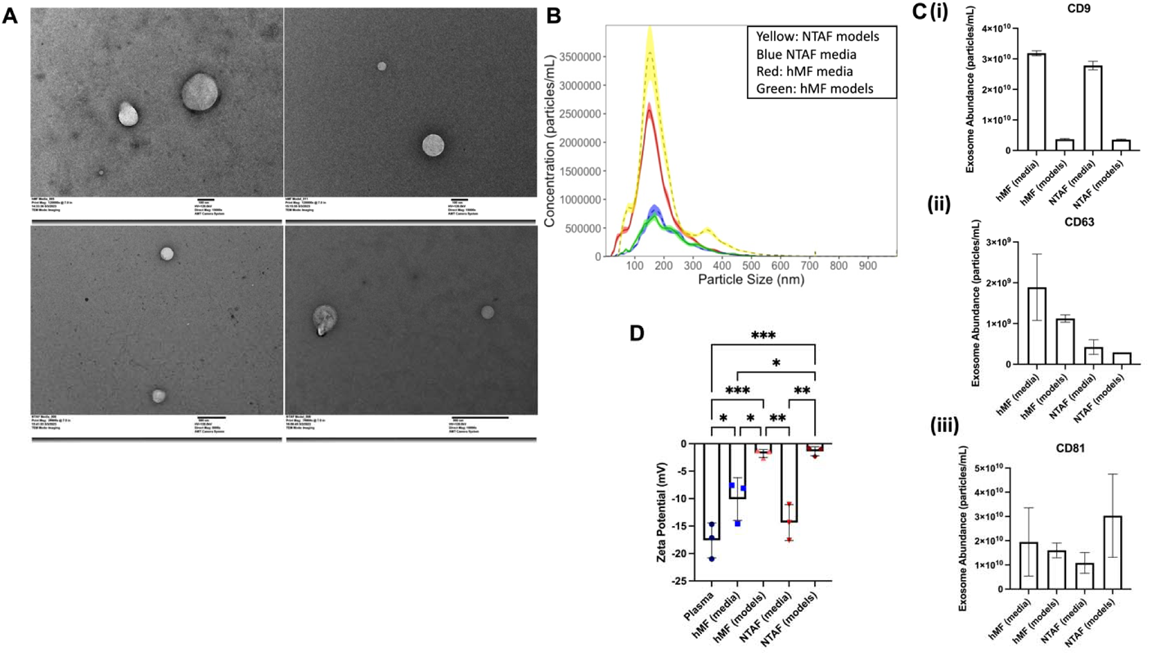
Characterization of EV groups. A) TEM images of hMF media EVs (top left), hMF model EVs (top right), NTAF media EVs (bottom left), NTAF model EVs (bottom right). B) NTA results of size and concentration distribution for the EV groups (n = 5). C) Exo-ELISA results for (i) CD9, (ii) CD63, (iii) CD81, exosome surface markers. D) Zeta potential results for the model EV groups and plasma control group. Data presented as the mean ± SD. ANOVA followed by Tukey’s post hoc was applied for statistical significance. *p<0.05, **p<0.01, and ***p<0.001.

### Cancer progression related cytokines are increased in the NTAF media group

Finally, we characterized the cytokines within the EVs from the different groups (Figure 4). Overall, there are six cytokines that are upregulated in hMF media group with the rest of the cytokines having lower regulation. The NTAF media group has consistent upregulation through most of the cytokines present in the array as compared to the rest of the groups. The cytokine array results show 35 cancer promoting cytokines that have increased expression in the NTAF media group. In contrast, the hMF media group showed increased expression for 13 cytokines that support cancer progression.

**Figure 4:**
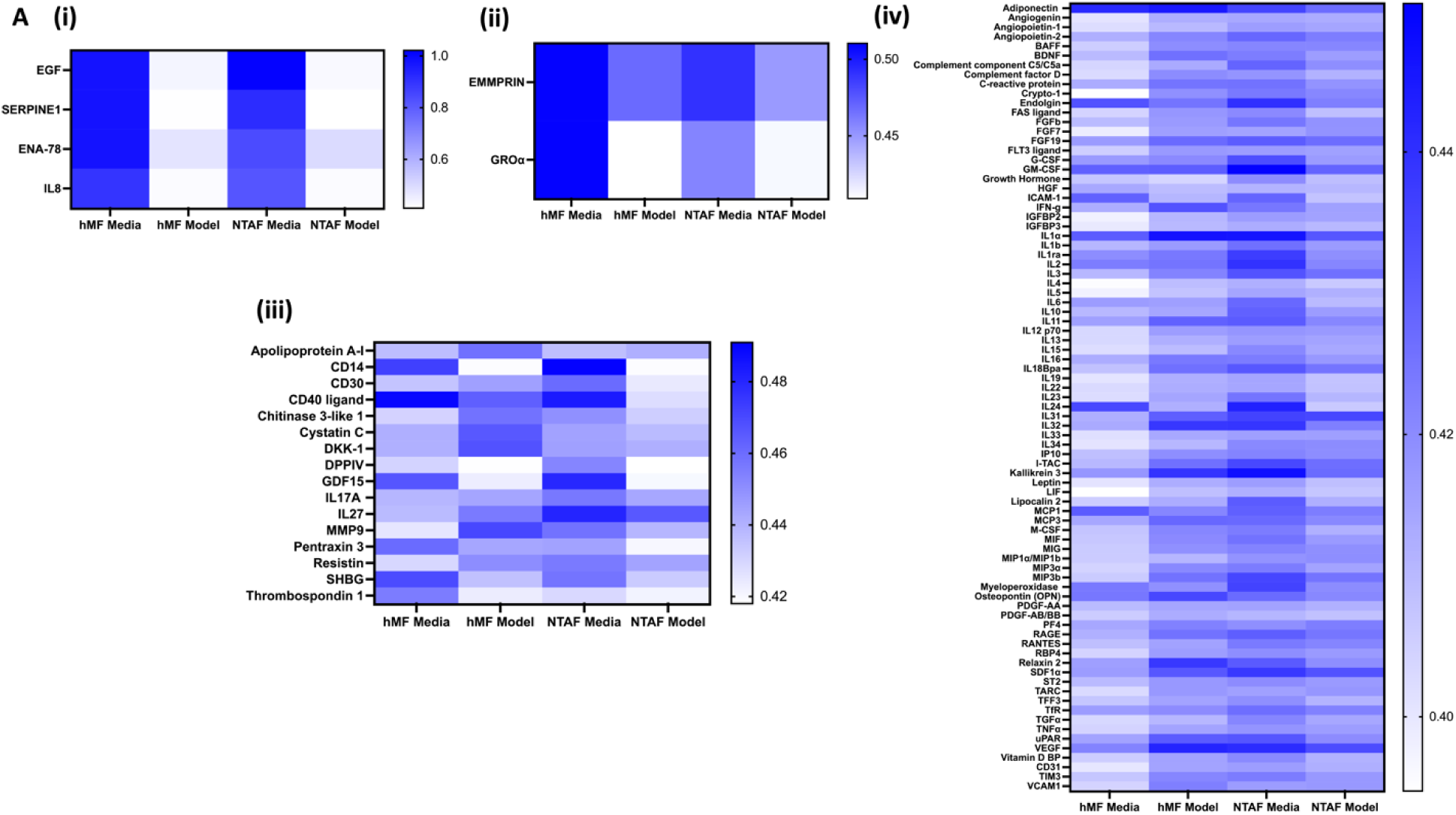
NTAF media has increase in cancer progression related cytokines. A) (i) Top four cytokines expressed in the cytokine array. (ii) Middle two cytokines expressed across the four model groups. (iii) Lower middle expressed cytokines in the cytokine array. (iv) Lowest expressed cytokines.

### Breast cancer cell migration, invasion, and proliferation is increased with NTAF EV treatment

2D cell migration assays were completed to test the effects of the EV groups on breast cancer cell behavior (Figure 5A). MDA-MB-231 cells were seeded into a 12-well plate and allowed to attach before treatment with the EV groups. Cells were treated with 20 µg of EVs for 48 hours. After treatment, cells were seeded in glass-bottom dish. Migration assays were conducted by imaging the cells seeded on a glass-bottom dish for 15 minutes every 10 hours. The cells were then tracked and the motility calculated. It was found that the NTAF groups had increased motility as compared to the hMF groups. NTAF media EVs caused the most significant increase in motility of 4.12 ± 0.82 µm/hr (Figure 5B). These results show that the NTAF group EVs cause an increase in breast cancer cell progression.

**Figure 5:**
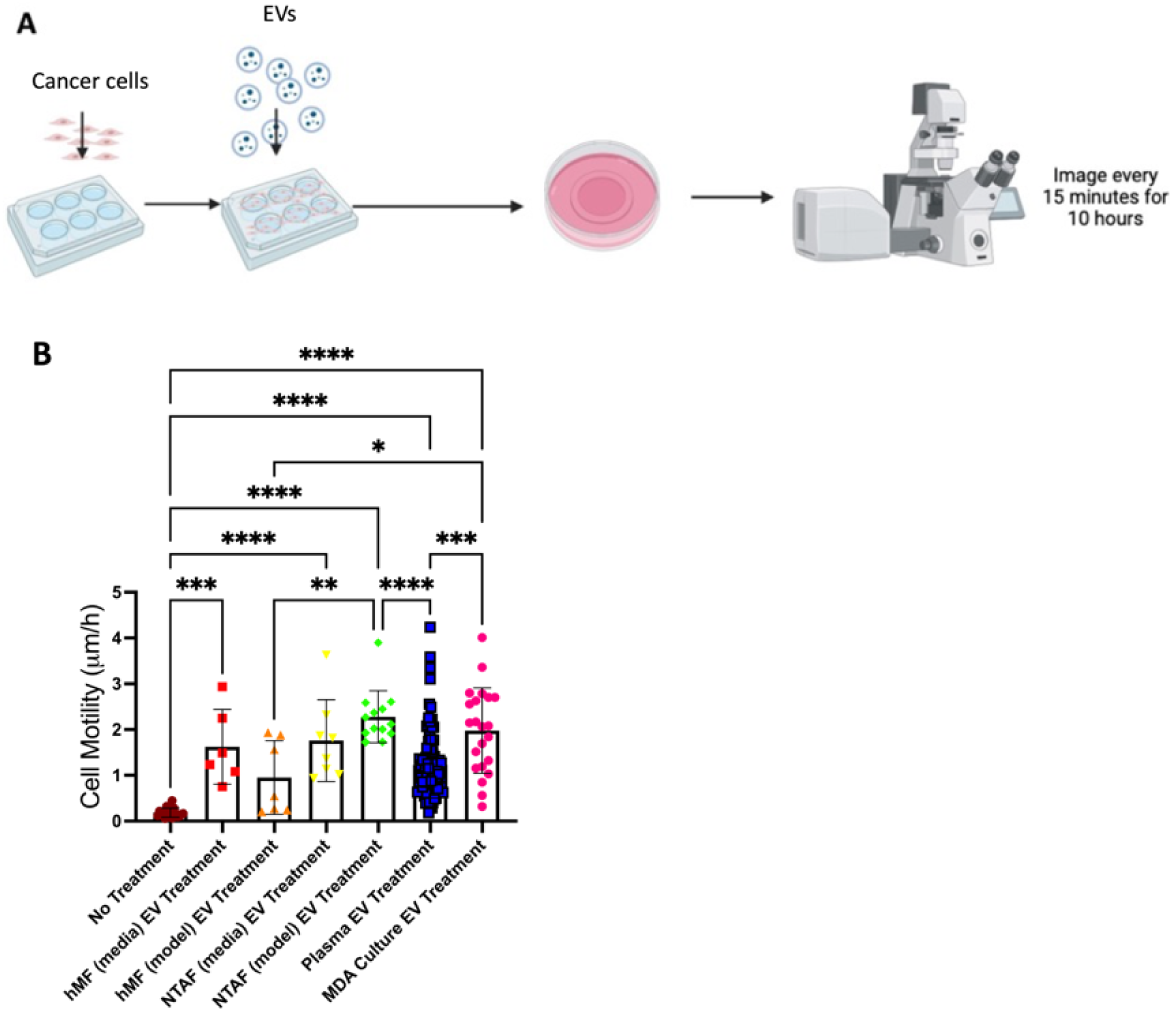
Cell motility in breast cancer cells is increased when treated with NTAF EV groups. A) Migration assay schematic. B) Cell motility assay results for each of the EV group treatments, no treatment control group, plasma EV treatment control group, and breast cancer cell culture EV treatment control group. Data presented as the mean ± SD. ANOVA followed by Tukey’s post hoc was applied for statistical significance. *p<0.05, **p<0.01, ***p<0.001, and ****p<0.0001. N = 3

**Figure 6:**
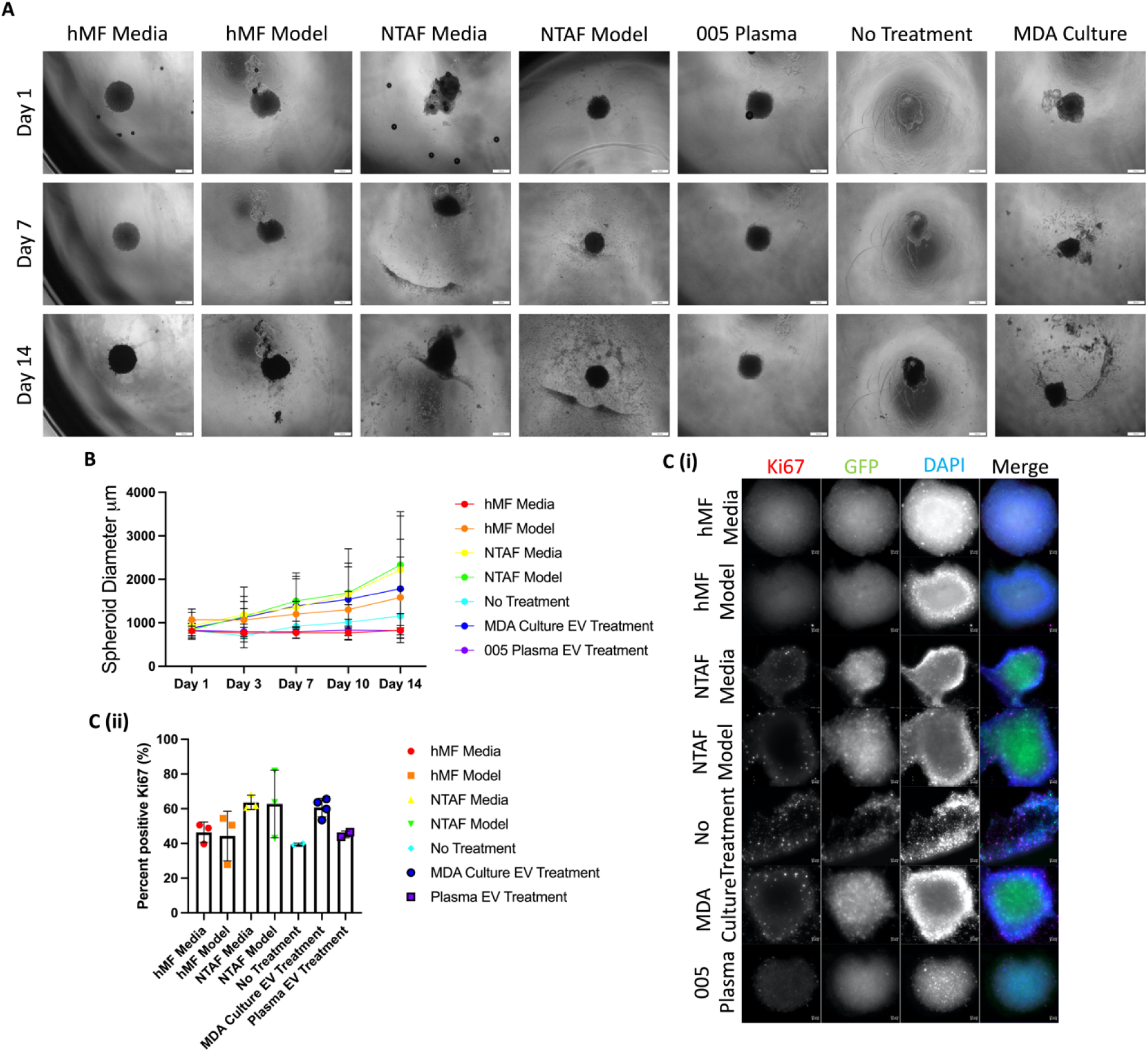
Local invasion is increased in a 3D model of breast cancer when treated with NTAF group EVs. (A) 3D breast tumor spheroid models brightfield images over the 14-day culture period. Scale bar: 500 μm. (B) Spheroid diameter quantification 3D breast tumor spheroid models. (C) (i) Ki67 staining of 3D models after 14 day culture period. Scale bars: 50 μm. (ii) Quantification of Ki67 stain. N = 2.

### Treatment with NTAF EVs increases tumor spheroid proliferation and invasion

Next, we utilized our tumor spheroids to create 3D breast tumor models. After model creation, these were treated with the EVs from the four EV groups. The breast tumor models were cultured for two weeks with images taken every three days to track spheroid diameter (Fig. 8A). The 3D models treated with the NTAF media EVs had the largest spheroid diameter increase of 1800 µm. The models were then fixed and stained for Ki67 (Fig. 8C(i)) showing that this invasion was due to the proliferation of the tumor spheroid. The 3D model treated with NTAF media EVs showed the highest percentage of cells positive for Ki67 at 63.7% ± 4.1%. Overall, these results suggest that the EVs secreted from normal tumor adjacent fibroblasts are fueling breast cancer progression.

## Discussion

This study was able to show distinct differences in the EV populations from fibroblasts of differing proximities to a breast tumor. This is the first such study to investigate the effect of distance from a tumor on fibroblast secretions using 3D bioprinted constructs to obtain EVs. We printed fibroblasts in a collagen gel to model the stromal compartment of the breast. The models included only fibroblasts in order to discern the effects of the different fibroblasts without any confounding factors. EVs were isolated from the effluent media from the models and from the models themselves after a week in culture. These EVs showed average exosome size and morphology. The hMF models produced higher concentrations of EVs in the media and lower concentrations within the models themselves. The opposite was seen in the NTAF models with higher concentrations of EVs within the models as compared to the media EV concentration.

Through ELISA, we were able to quantify the classic exosome surface markers CD9, CD63, and CD81. The different groups showed different zeta potentials with the EVs isolated from the models having a zeta potential closer to zero. Cytokines within the EVs were analyzed using a dot-based assay and found individual cytokine profiles for the different groups. The NTAF media group showed an upregulation of cytokines related to breast cancer progression. After this, we treated breast cancer cells with the different EV groups to study how they affected the resulting breast cancer cells migration. The breast cancer cells treated with the EVs from the two NTAF groups showed increased migration, invasion, and proliferation with the largest increase in these three categories stemming from treatment with the NTAF media group. A 3D spheroid model was then used to investigate the effect of these EVs in a more biologically relevant system. Our results showed an increase in local invasion and proliferation for the spheroids treated with the NTAF media group. Through this investigation we were able to prove that normal fibroblasts that reside next to a tumor secrete EVs that support breast cancer progression regardless of EV origin.

Once the EVs were isolated from the media and the models, we characterized them using TEM, NTA, ExoELISA, and zeta potential measurements. TEM and NTA showed an average particle size of 100 nm to 200 nm, which is within the range of average exosome sizes ^53–55^. ExoELISA confirmed the presence of traditional exosome surface markers of similar concentrations for CD63 and CD81. Our results also showed a decrease in CD9 for the EVs isolated from the models. This is comparable with matrix bound vesicles (MBVs) as they have been shown to have decreased CD9 as well ^14,16^. NTA showed increased concentrations of EVs within the NTAF models. The effluent media from the NTAF models had a lower concentration as compared to the models. We see the opposite result for the hMF models with increased concentration within the media as compared to the models. This was not the case for direct comparison between liquid-phase EVs and MBVs ^16^, so this could be due to surface charge differences between the EVs from the groups. In order to assess if this change in concentration was due to differences in surface charges on the EVs, we then measured the zeta potential of each of the EV groups.

Multiple studies ^56–61^ have characterized the zeta potential of conditioned media from cells, human plasma, and various other EV sources and found zeta potentials in the range from 0 mV to -50 mV, putting our measurements within the reported range. EVs isolated from effluent media of diseased cells were shown to have decreased zeta potential as compared to EVs from healthy cells ^57,59,61^, which is similar to our results showing decreased zeta potential of the NTAF media EVs as compared to the hMF media EVs. The model groups showed zeta potentials closer to zero, and as collagen is negatively charged ^62,63^, this could explain one method for the model EVs to attach to the collagen within the model as the more negatively charged media EVs would be repelled by the collagen and stay within the media. However, this conflicts with the results we see from our mouse breast EVs which show a significantly decreased zeta potential as compared to the other groups. This could be due to many other factors within the breast tissue such as differing ECM proteins, cell types, and cytokines present that could affect the zeta potential of this group of EVs. This difference in zeta potential could explain the difference in concentrations between the NTAF model and NTAF media EVs. Surface characterizations of the EVs showed standard EV surface proteins, size, and shape. The next characterizations were performed on the cargo of the EVs.

The cytokine array showed different cytokines upregulated within the cargo of the EV groups. Visually, our results indicate that the hMF media group has some cytokines that are highly expressed, whereas the other three groups have more expression across the entirety of the panel. Our results are comparable to other labs comparing matrix bound vesicles (MBVs) to liquid-phase EVs as other studies also showed more proteins within the liquid-phase EV cargo as compared to the MBV cargo ^16^. Based on a study’s ^16^ silver stain of electrophoretically separated proteins, the stain for the MBVs was visually lighter than the liquid-phase EVs, suggesting less proteins in their cargo. Multiple studies ^16,64^ also found differences within the miRNA content between these two groups. Phospholipids were found to be significantly higher within the MBVs, which could explain the difference in cargo and the decreased cytokine content within our model group EVs. The MBVs also showed a significant increase in fatty acid cargo as compared to the liquid-phase EVs, with the group determining that the MBV cargo is a pool of substrates for the synthesis of signaling lipid mediators. This could mean that there are more phospholipids, RNAs, or miRNAs within the cargo of the MBVs rather than cytokines.

Of the cytokines with increased expression within the NTAF media group, epidermal growth factor (EGF) is one cytokine of interest in our results. EGF has consistently been shown to increase proliferation in breast cancer cells ^65–72^. One study also showed that after treatment with EGF, breast cancer cells showed increased cell speed and directionality and that cells migrated towards a higher EGF concentration ^71^. Another cytokine of interest, IL-27, is shown to be increased in NTAF groups and specifically in the NTAF media group and the lowest level in the hMF media group. Clinically, IL-27 was increased in serum of breast cancer patients as opposed to the control ^73^. IL-27 has been shown to have dual roles tumor immunity through both working with IL-12 to promote tumor self-recognition with T-helper cells to allow the tumor to avoid the immune system and inducing cytotoxic T-lymphocytes which are anti-tumor ^74–76^. Overall, IL-27 can either promote anti-tumor immunity or support tumor progression. CD-14 was increased in the media groups and more-so in the NTAF media group. CD-14 is typically a marker of macrophages and increased CD-14 positive macrophages within the breast primary tumor has been shown to have increased cancer relapse ^77^ while increased CD-14 cytokine content in serum is correlated with increased breast cancer recurrence ^78^. CD-14 has also been implicated in tumor-promoting inflammation, proliferation, and overall tumor growth ^79,80^. Finally, GDF15 is most highly expressed in the NTAF media group. GDF15 has been shown to support and even be necessary for chemotherapeutic and radiation resistance in breast cancer cells ^81–84^. GDF15 has also been shown to be required for breast cancer cell growth ^84^, cancer stem cell maintenance ^85^, and is repressed by YAP ^86^. These cytokines that are upregulated in the NTAF groups can influence the behavior of breast cancer cells and have an effect on the progression of breast cancer.

We next performed cell-based assays to investigate the effect of the different EV groups on breast cancer cell behavior. Overall, we saw an increase in the migration, invasion, and proliferation, or EMT-like behavior, for breast cancer cells treated with the NTAF groups, with the largest increase coming from those treated with the NTAF media EVs. The plasma EV treated cells showed an average migration, invasion, and proliferation similar to those treated with the normal fibroblast EV group as expected as treatment with EVs increases the aggressiveness of cells ^87–89^. The MDA-MB-231 effluent culture media EVs caused a large increase in this progressive nature of the cells, which is supported in literature as breast cancer cells secrete EVs that support breast cancer cell EMT-like behavior ^11,90,91^. The NTAFs’ EV secretion is being influenced by its proximity to the tumor. CAFs have been previously shown to increase breast cancer cell migration ^92–94^, invasion ^95–97^, and proliferation ^96,98,99^ and their EVs contain cytokines that promote breast cancer progression ^20–22^. NTAF secreted EVs are similar to the CAF EVs in support of breast cancer progression while the hMF EVs have decreased support as compared to these populations.

Finally, we utilized our 3D breast tumor model to study spheroid progression as a result of EV group treatment. The NTAF media EV group treated spheroids had the largest local invasion as demonstrated by the increase in diameter and the most increased Ki67 positive stain. In the cell-based assays we see increased EMT-like behavior for cells treated with NTAF group EVs regardless of EV localization. There are demonstrated differences between the EVs of different origin such as the concentration or zeta potential. These differences likely affect the localization of the EVs, but not as strongly as the proximity to a tumor affects these EVs. The fibroblast phenotype overrides the EV location differences in resulting breast cancer cell behavior due to EV treatment. Here we show that the NTAF secretome is being influenced by the tumor and is affecting the tumor that they surround. Again, this aligns with previous literature that demonstrates that CAFs promote breast cancer cell invasion ^95–97^ and proliferation ^96,98,99^, showing that these NTAF cells are more similar to and have influences from CAFs. Our study suggests that NTAFs are part of a gradient of fibroblasts phenotypes based on proximity to a tumor rather than having only normal fibroblasts and cancer-associated fibroblasts. Other studies have shown that stromal cells such as fibroblasts or mesenchymal stem cells have an effect on a 3D tumor spheroid through proliferation and invasion of the tumor spheroid ^100–103^, further confirming our findings. Overall, our results suggest that NTAF cells have an altered secretome from that of normal fibroblasts, regardless of EV secretion location, and the EVs from the secretome aid in breast cancer progression.

## Conclusion

In this study, we show that the NTAF cells produce EVs that contain cytokines that promote breast cancer progression and influence the behavior of breast cancer cells despite differences in EV secretion location. These cells have a distinct EV secretome different from normal human mammary fibroblasts, showing that the proximity to a tumor has an effect on the fibroblasts surrounding the tumor. While there were differences between the EVs from the media groups compared to those of the model groups, these differences did not have as much of an effect on the resulting breast cancer cell behavior as the fibroblast phenotype. This can be investigated in further studies along with factors that are increased in areas surrounding the tumor which help the tumor progress and give researchers targets for treatment to slow this progression. The 3D bioprinted breast tumor model created in this study can also be used in the future to test novel breast cancer drugs, decrease the usage of animal studies, and test drugs on a more human replicative environment. By 3D bioprinting our model, this can allow future researchers to have a high throughput system to study breast cancer progression.

## Author Contributions

Conceptualization: Pinar Zorlutuna, Jensen N. Amens

Data curation: Jensen N. Amens, Jun Yang, Lauren Hawthorne Formal analysis: Jensen N. Amens

Funding acquisition: Pinar Zorlutuna, Jensen N. Amens Investigation: Jensen N. Amens, Jun Yang, Lauren Hawthorne Methodology: Pinar Zorlutuna, Jensen N. Amens

Project administration: Pinar Zorlutuna Resources: Pinar Zorlutuna

Software: Jensen N. Amens Supervision: Pinar Zorlutuna Validation: Jensen N. Amens Visualization: Jensen N. Amens

Roles/Writing - original draft: Jensen N. Amens

Writing - review & editing: Jensen N. Amens, Jun Yang, Lauren Hawthorne, Pinar Zorlutuna

## Data Availability Statement

Data will be made available upon request.

## Acknowledgments

This study was supported by The Interdisciplinary Interface Training Program (IITP) Grant through the Walther Cancer Foundation. This study was funded by NIH award number R01 CA275423-01A1. We would also like to acknowledge Dr. Siyuan Zhang for the gift of the MDA-MB-231 cells used in this study and Dr. Harikrishna Nakshatri of Indiana University for the gift of the fibroblasts used in this study. This work is partially supported by the Notre Dame Integrated Imaging Facility.

